# Evidence of divergent selection in a parasite due to its host immunological respond

**DOI:** 10.1101/2022.05.15.492026

**Authors:** Kum C. Shim, Jesse N. Weber, Stijn den Haan, Daniel I. Bolnick

## Abstract

We investigated if an immunological response (i.e. fibrosis) in threespine stickleback fish can cause divergent selection in its tapeworm *Schistocephalus solidus*. We pooled tapeworms from six populations in Vancouver Island (BC, Canada) and sequenced the whole genome of these pools. Then we used a modified Population Branch Statistics (PBS) technique based on F_ST_ comparisons to find loci under divergent selection due to fibrosis. We found at least eight loci under divergent selection in tapeworm populations caused by this strong immunological respond to infection from the fish.

## Introduction

There is little doubt that parasites are a strong selective force to their hosts (Weber et al. 2021, Fumagalli et al. 2011, Ebert and Fields 2020). The opposite is also true: hosts can also be a major selecting force in the evolution of parasites (Schmid-Hempel 2008, Maizels et al. 2004). The product of such coevolution has not only been linked to the maintenance of genetic diversity between and within species (Ebert and Fields 2020) but also to major evolutionary transitions such as haploidy to diploidy, selfing to outcrossing, and the evolution of sex (Nuismer and Otto 2004, Becks and Agrawal 2012, Hamilton 1980, Lively 1987, Morran et al. 2011). Thus, understanding coevolution at the genetic level should be of major importance in evolutionary studies and should provide insights into these major evolutionary transitions.

Most of the genetic work on host-parasite coevolution, especially for macroparasites like helminths, has been focused on the host’s adaptation to parasites (Weber et al. 2021, Nuismer et al. 2017, Atlija et al. 2016, Benavides et al. 2015, Kim et al. 2015, Wilfert and Schmid-Hempel 2008, Kover and Caicedo 2001), but to understand the genetic details of coevolution and how genes interact between hosts and parasites, one must also focus on the parasites. Here we studied the genetic basis of coevolution from the parasite’s perspective, by assessing genomic signatures of divergent selection acting on a helminth, spanning host populations with divergent immune traits. For this we used the threespine stickleback-*Schistocephalus solidus* tapeworm system.

The threespine stickleback (*Gasterosteus aculeatus*) is a small fresh water and marine fish found in the Holarctic. In freshwater, they are parasitized by the tapeworm *Schistocephalus solidus* (Dubinina 1980). The tapeworm is very host specific to the stickleback and can reach high infection prevalence in some fish populations (Weber et al. 2021). High infection loads can be detrimental to the fish due to increased internal inflammation, behavioral changes leading to higher predation rates, and lowering reproductive output (Weber et al. 2021, De Lisle and Bolnick 2020, Talarico et al. 2017); consequently, the sticklebacks present a wide range of behavioral, physiological, and immunological mechanisms to fight tapeworm infections (Weber et al. 2021, Fuess et al. 2020, Weber et al. 2017).

Recent research suggests that fibrosis is a key element in sticklebacks’ ability to successfully limit *S. solidus* infection. Peritoneal fibrosis (or simply fibrosis) in sticklebacks is the buildup of connective tissue like collagen and fibroblast within the peritoneal cavity. The peritoneum is a single cell-thick layer of tissue lining the body cavity in vertebrates. Tapeworm-infected sticklebacks develop fibrosis that form a cocoon around the major organs within the peritoneal cavity, and the organs may even adhere to the peritoneum. The processes in which fibrosis combats helminth infections in sticklebacks start by first curtailing the growth of the tapeworms in its body cavity, and second by enveloping and killing the helminth (Weber et al. 2021). Fibrosis is a heritable response in sticklebacks to parasite infections (Weber et al. 2021, Hund et al. 2020), and it is caused by the inflammation response to tapeworm migration from the fish gut to the body cavity (Weber et al. 2021; Mittal et al. 2014). Tapeworm migration through the gut wall happens 12-24hr after tapeworm infection in the fish (Hemmerschmit and Kurtz 2007). The fibrosis adhesions can have a negative effect on the sticklebacks such as decreasing female fitness and affecting male’s nesting care (Weber et al. 2021, De Lisle and Bolnick 2020). Due to this combination of benefits and costs of fibrosis, some lake populations of stickleback exhibit extensive fibrosis, while others have apparently evolved secondary loss of fibrosis to mitigate costs, instead adopting a tolerant strategy (Weber et al. 2021).

Past field surveys have shown that stickleback populations with high levels of fibrosis tend to have on average smaller tapeworms (Weber et al. 2021). However, there is variance around this negative relationship between fibrosis and cestode size. In past field surveys, we noted that in some lakes the native tapeworms are able to grow comparatively large despite a high incidence of severe fibrosis in their local stickleback hosts (table 1). Thus, are there parasite loci that are consistently divergent between high fibrosis versus low fibrosis fish lakes? This question is important because these loci could be the start of a chemical and/or physiological pathway to allow some tapeworm populations to adapted and overcome fibrosis in their local sticklebacks.

**Table 1.** Sample locations, dates, sample sizes, and fibrosis status:

| lake | latitude | longitude | collection date | watershed | Fish N<sup>1</sup> | Tapeworm N<sup>2</sup> | Fibrosis rate (%)<sup>3</sup> | Infection rate (%)<sup>4</sup> | Average tapeworm size (g) | Fibrosis status |
| --- | --- | --- | --- | --- | --- | --- | --- | --- | --- | --- |
| Lawson | 50.035596 | -125.57831 | 6/24/16 | Campbell River | 31 | 21 | 51.61 | 29.03 | 32.15 | High Fibrosis |
| Boot | 50.055025 | -125.52644 | 6/24/16 | Campbell River | 31 | 23 | 74.19 | 87.1 | 35.19 | High Fibrosis |
| Gosling | 50.045922 | -125.5014 | 6/24/16 | Campbell River | 31 | 24 | 29.03<sup>5</sup> | 41.94 | 17.46 | Low Fibrosis |
| Echo | 49.987652 | -125.41133 | 6/19/16 | Campbell River | 41 | 25 | 2.78 | 43.9 | 62.71 | Low Fibrosis |
| Frederick | 48.858001 | -125.02306 | 6/29/16 | Alberni Inlet | 30 | 24 | 40 | 80 | 11.33 | High Fibrosis |
| Pachena | 48.837661 | -125.03389 | 7/2/16 | Alberni Inlet | 7 | 23 | 0 | 71.43 | 26.10 | Low Fibrosis |

<sup>1</sup> Number of threespine stickleback fish collected for this work.
<sup>2</sup> Number of *Schistocephalus solidus* tapeworms collected per lake for this work from the previous fish N column. This number also represent the number of tapeworms pooled per lake.
<sup>3</sup> Fibrosis rate in the threespine stickleback fish from each lake.
<sup>4</sup> Tapeworm infection rate in the threespine stickleback fish from each lake.
<sup>5</sup> In surveys from other projects, the fibrosis rate for Gosling Lake sticklebacks has been documented to be as low as 6% (Hund et al. 2020)

To answer the above question, we used whole-genome Pool-seq (Schlötterer et al. 2014) of six tapeworm populations, three from fish populations with high incidences of fibrosis and three from populations with lower fibrosis frequencies. Then, we compared the genomes from both population sets to find regions of extreme genetic differences, which would suggest an effect of divergent natural selection between closely related tapeworm populations. Finally, we tested for tapeworm loci that are consistently divergent between high versus low fibrosis populations of stickleback. Indeed, we found that even though the six tapeworms populations tested here were not genetically very distinctive, they still had a few loci that were under strong divergent selection, and at least eight of these loci to be consistently divergent between the high versus low fibrosis fish populations.

## Materials and Methods

### Tapeworm collecting

The *S. solidus* tapeworms were collected in June and July 2016. For more details on the location and sample sizes, see table 1 and figure 1. The threespine stickleback fish were collected using un-baited minnow traps left submerged overnight in shallow water (< 3m) along each lake’s shorelines (Scientific Fish Collection Permit No. NA16-230545, Ministry of Forest, Lands and Natural Resource Operations, BC, Canada). A random sample of captured fish were euthanized in MS-222 and dissected on-site (methods approved by The University of Texas Institutional Animal Care and Use Committee as part of AUP-2010-00024). The size (i.e. weight and length), sex, tapeworm infection status, and fibrosis of each fish were recorded during dissection. The tapeworms were stored in 1.5mL centrifuge tubes (one per fish) and preserved in 100% ethanol for later DNA extraction.

**Figure 1:**
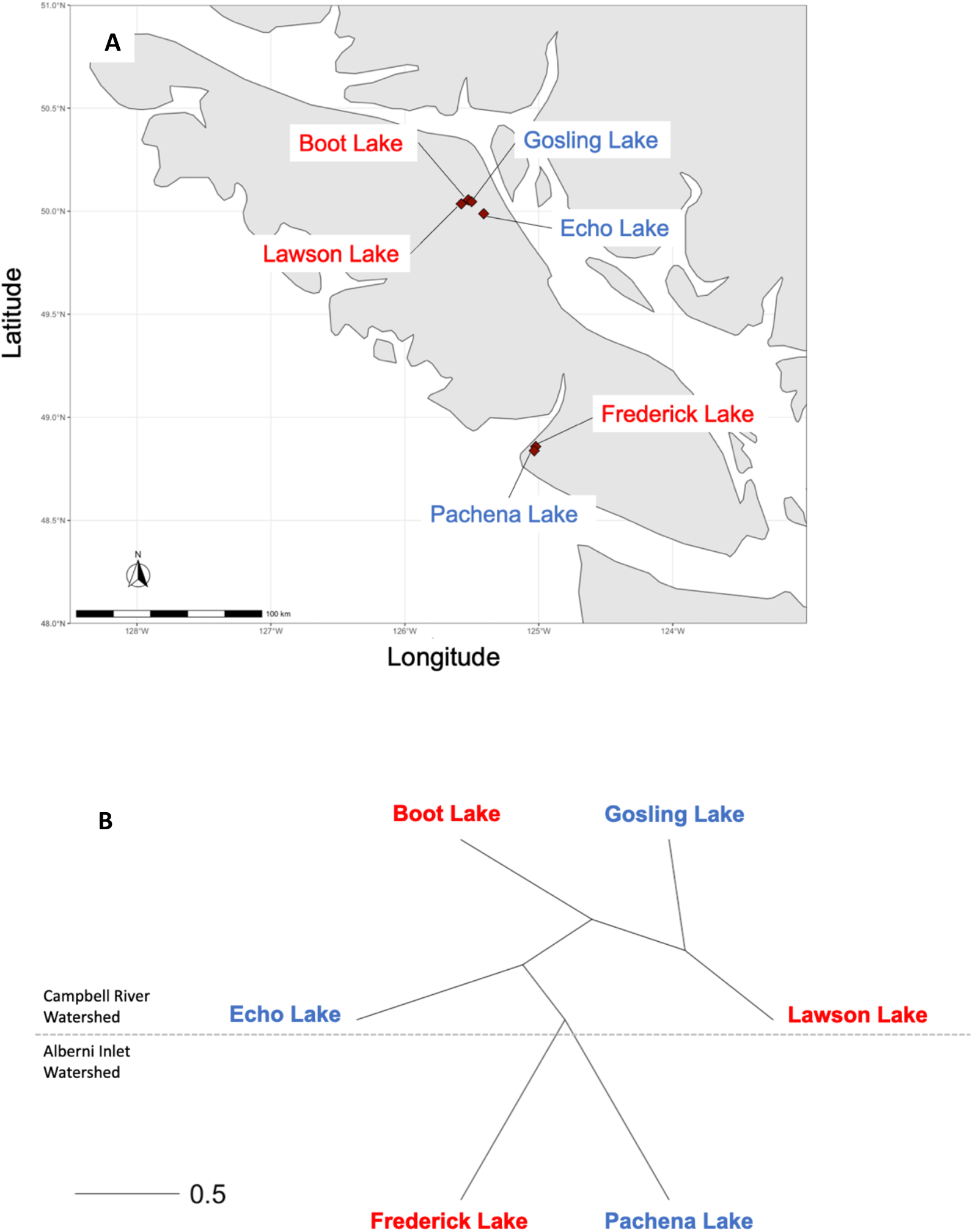
Lake locations in Vancouver Island (A) along with a Neighbor Joining tree (from genome-wide F_ST_ values) for the tapeworm populations from each lake (B). High fibrosis lakes are colored red and low fibrosis lakes are colored blue. The dash line in the Neighbor Joining tree separates the lakes from Campbell River Watershed (those above the line) from the Port Alberni Inlet lakes (those below the line).

### DNA extraction and pooling

The DNA for each tapeworm were extracted back in our laboratory in Texas using the Wizard® SV 96 Genomic DNA Purification Kit and System by Promega (Madison, WI).

The DNA concentration was quantified using Quan-iT PicoGreen dsDNA Assay Kits and dsDNA Reagents by Thermofisher (Waltham, MA) and a 96-well microplate reader (Tecan Infinite M200 Pro, Mannedorf, Switzerland)

After quantification, a specific amount of DNA (see supplementary table 1) was collected from each tapeworm and pooled per lake. Whole-genome Pool Sequencing is a very cost-effective method of comparing the genome-wide allele frequency differences between populations. As long as the pooling is from a large number of individuals from each population, the allele frequency estimates from pool sequencing are accurate and reliable, and it is effective in SNP discovery (Schlötterer et al. 2014). The number of tapeworms pooled per lake is shown in table 1. The pooled DNA was sent to Genohub (Austin, TX) for paired-end next generation whole-genome sequencing using an Illumina HiSeq sequencer. The read lengths were 2 × 150bp.

### Bioinformatics

The raw sequences were trimmed from the Illumina adaptors using bbduk.sh from BBMap bioinformatic tools version 38.62 (Bushnell et al., 2017). Sequences were then mapped to the *S. solidus* tapeworm reference genome from the Aubin-Horth group, Université Laval (Berger et al., 2021) using Samtools version 1.6 (Danecek et al., 2021). The sequences were then filtered for a minimum mapping quality of 20 and then joined into one mpileup file, both using Samtools. The mpileup file was then converted into a “synchronized” file using PoPoolation2 (Kofler et al., 2011). This synchronized file was then uploaded into Poolfstat version 1.1.1 (Hivert et al., 2018), a R package (R Core Team) to compute pairwise F_ST_ between the pooled populations, for each SNP and each pairwise combination of populations. Negative F_ST_ for any SNP found by Poolfstat were replaced by zero.

### Finding possible SNPs under selection due to fibrosis in sticklebacks

To identify loci under divergent selection between the tapeworm populations due to fibrosis in fish, we used the following calculation:

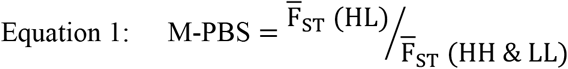

Where:

**M-PBS**: Multipopulational PBS

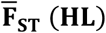: Mean F_ST_ for High vs Low Fibrosis Lakes comparisons (see table 1 for High and Low Fibrosis lakes)

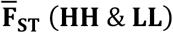: Mean F_ST_ for High vs High and Low vs Low Fibrosis Lakes comparisons

Equation 1 is a modification of Population Branch Statistics (PBS; Yi et al., 2010) which was originally developed to identify targets of selection in threeway comparisons between populations (an outgroup and two divergent populations). Mark Kirkpatrick (pers. comm) adapted this PBS to reflect genetic distances among two sets of populations from different environments or with different traits (i.e. Multipopulational PBS).

To avoid possible false positive results due to chance, we randomly permuted the fibrosis status of each lake (but keeping three high and three low fibrosis lakes) and repeated calculations from equation 1. Because two populations are geographically distant (Frederick and Pachena lakes), we only used permutations that retained one high and one low fibrosis assignment in this region. We then compared the observed peaks of Multipopulational PBS, against the values obtained via permutation, to evaluate whether stronger PBS is observed between high and low fibrotic populations. Because of the limited permutations of populations, this does not reflect a formal permutation-based p-value.

We also made Manhattan plots of 500 SNP windows for all F_ST_ pairwise comparisons in R (R Core Team). This is to visualize F_ST_ peaks (if any) for all F_ST_ comparisons; these peaks might indicate SNPs under divergent selection for the populations being compared irrespective of fibrosis status.

## Results

We recovered 4,901,696 SNPs from the whole genome pool sequencing of the six *S. solidus* tapeworm populations. The six population pools were sequenced to an average depth coverage of ∼334 reads per site. The average number of reads per site per pool range from 29.61 to 46.39 (see supplementary table 4 for each population pool’s average number of reads). The tapeworm genome size is ∼540Mbp in size (WormBase in Howe et al. 2017). The genome-wide F_ST_ between the tapeworm populations range from 0.0 to 0.02, and the genome-wide heterozygosity for the populations averaged to 0.001 (see table 2). The neighbor joining tree based on genome-wide F_ST_ clustered the tapeworm populations according to watershed and geographical distribution (see figure 1).

**Table 2.** Genome-wide F_ST_ and heterozygosity for the six tapeworm populations.

|  | Boot lake | Echo lake | Frederick lake | Gosling lake | Lawson Lake | Heterozygosity |
| --- | --- | --- | --- | --- | --- | --- |
| Boot lake |  |  |  |  |  | 0.0010 |
| Echo lake | 0.0026 |  |  |  |  | 0.0011 |
| Frederick lake | 0.0226 | 0.0214 |  |  |  | 0.0010 |
| Gosling lake | 0.0019 | 0.0032 | 0.0226 |  |  | 0.0010 |
| Lawson Lake | 0.0 | 0.0009 | 0.0211 | 3.44E-05 |  | 0.0010 |
| Pachena lake | 0.0171 | 0.0157 | 0.0018 | 0.0172 | 0.0158 | 0.0011 |

The Manhattan plots of 500 SNP windows for all F_ST_ pairwise comparisons indicate there were outlier peaks for most (13 out of 15) population comparisons. The number of peaks ranges from 3.1 to at least 3 per comparison (see figures 2 and 3). A closer evaluation indicated that most of these F_ST_ peaks were from comparisons between Campbell River Watershed lakes (i.e. Boot, Echo, Gosling, and Lawson lakes) and Port Alberni Inlet lakes (Frederick and Pachena lakes). The remaining high F_ST_ peaks were between Echo and the remaining Campbell River Watershed lakes (Boot, Echo, and Lawson). We identified a total of 22 SNPs (some shared among populations) responsible for these outlier F_ST_ peaks; their F_ST_ values range from 0.72 to 1.0 (see supplementary table 2). These F_ST_ peaks are possibly due to geographic differentiation (or local adaptation) between the watersheds since Campbell River Watershed and Port Alberni Inlet are ∼130km apart (figure 1A), and Echo Lake is also 10-15km apart from the remaining Campbell River watershed lakes.

**Figure 2:**
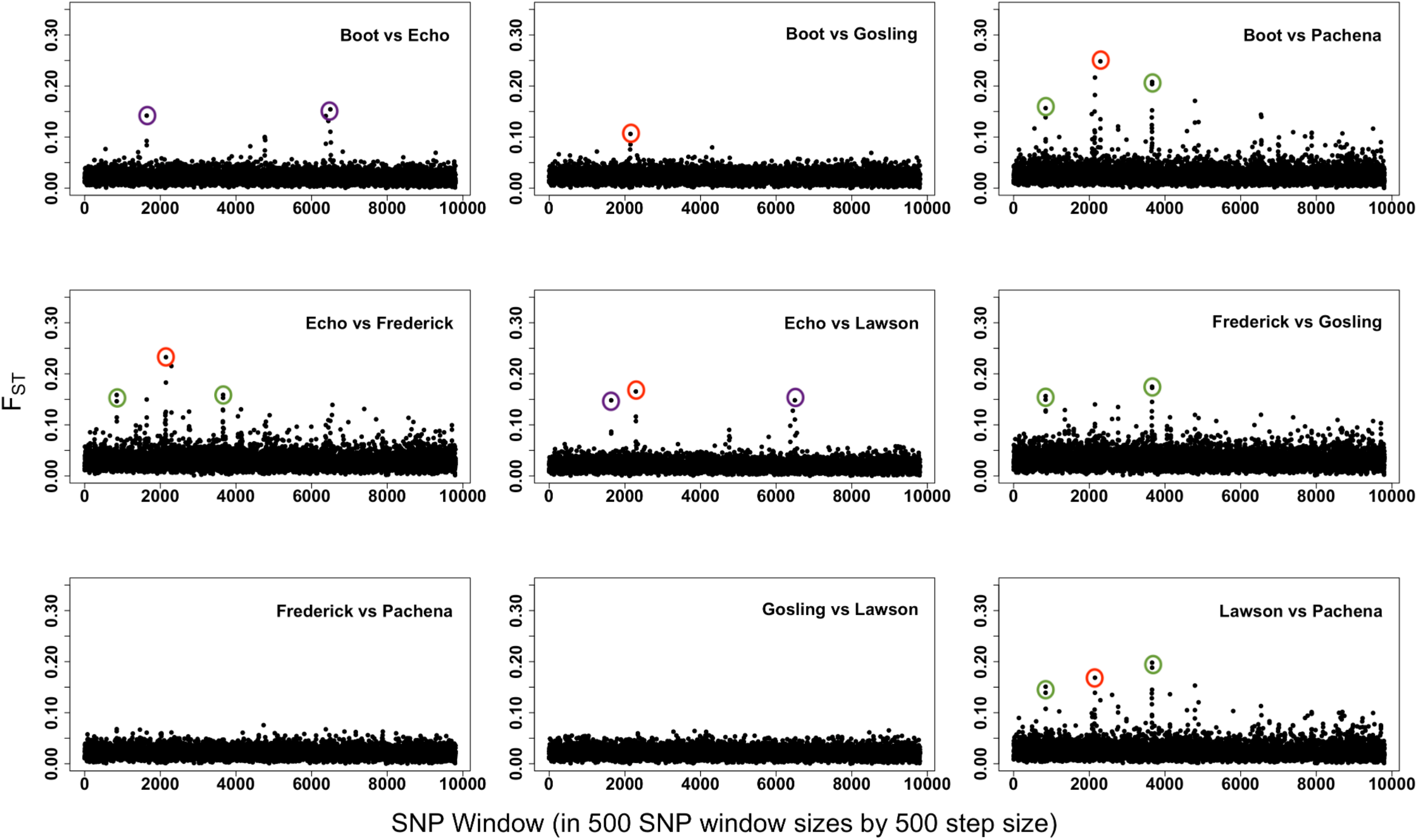
Manhattan plots of High vs Low (HL) Fibrosis Lake Comparisons (in tapeworms) for 500 SNP windows by 500 SNP step sizes with no discernible outlier F_ST_ peak shared by all comparisons. However, there were some outlier F_ST_ peaks shared by a few comparisons: around the 2000 SNP window (circled in red), in Campbell River vs Alberni Inlet Lake comparisons (circled in green), and Echo vs other Campbell River watershed Lake comparisons (circled in purple). These SNPs possibly signal local adaptations between watersheds and lakes. See supplementary table 2 for the exact location of these SNPs and their F_ST_ values. The labels inside each graph indicate each lake pair in the F_ST_ pairwise comparisons.

**Figure 3:**
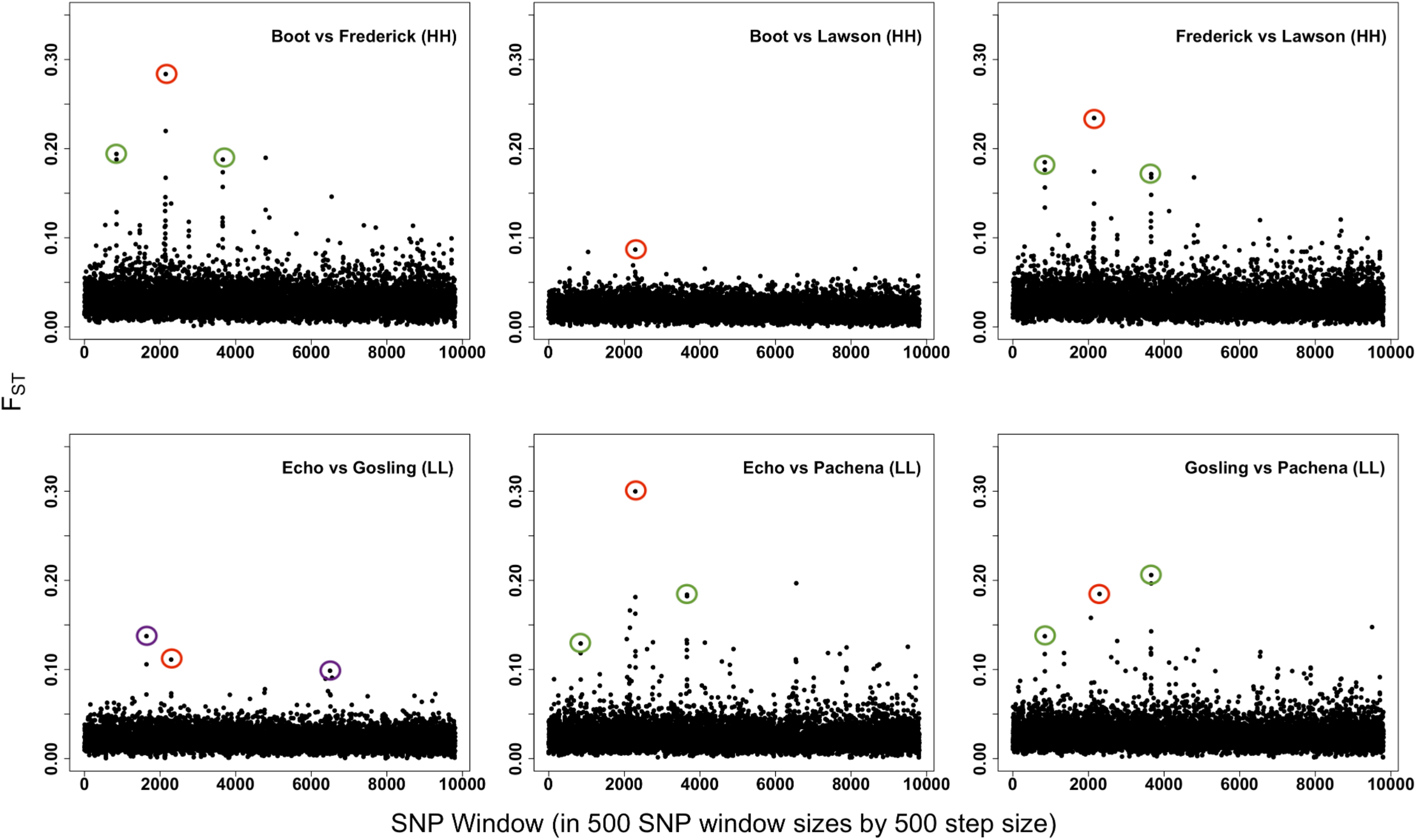
Manhattan plots of High vs High (**HH**) and Low vs Low (**LL**) Fibrosis Lake Comparisons (in tapeworms) for 500 SNP windows by 500 SNP step sizes. Like High vs Low Fibrosis Lake comparisons (figure 2) there were some outlier F_ST_ peaks shared by a few comparisons: around the 2000 SNP window (circled in red), in Campbell River vs Alberni Inlet Lake comparisons (circled in green), and Echo vs other Campbell River watershed Lake comparisons (circled in purple). These SNPs possibly signal local adaptations between watersheds and lakes. See supplementary table 2 for the exact location of these SNPs and their F_ST_ values. The labels inside each graph indicate each lake pairs in the F_ST_ pairwise comparisons.

When the Manhattan plots of 500 SNP windows were grouped into High vs Low (HL) Fibrosis Lake comparisons, they did not show consistent outlier F_ST_ peaks across all comparisons (figure 2). The same was true in High vs High (HH) and Low vs Low (LL) Fibrosis comparisons (figure 3). However, the M-PBS (i.e. equation 1) did show some SNPs with a high ratio, indicating relatively high F_ST_ only in the HL Fibrosis Lake comparisons (see figure 4 showing ratios for 500 SNP windows). We zoomed into the SNP windows to pinpoint the exact high ratio SNPs and found four potential SNPs corresponding to window #4762 and five potential SNPs for window # 6493 (see figure 5). After further assessment of the F_ST_ values of each of these SNPs, we kept all SNPs except #3246136 from window # 6493 for further analyses. These eight SNPs had consistent relatively high F_ST_ in the HL comparisons and zero or near zero F_ST_ in the HH and LL comparisons (see table 3 for all the F_ST_ values and the exact position of each of these SNPs in the reference genome). The remaining SNP #3246136 from window # 6493 had relatively low F_ST_ for HL and HH and LL comparisons (see table 3). Furthermore, we performed simple t-tests comparing the F_ST_ for HL vs HH and LL comparisons for each SNP, and SNP #3246136 from window # 6493 was the only non-significant among the SNPs (table 3).

**Figure 4:**
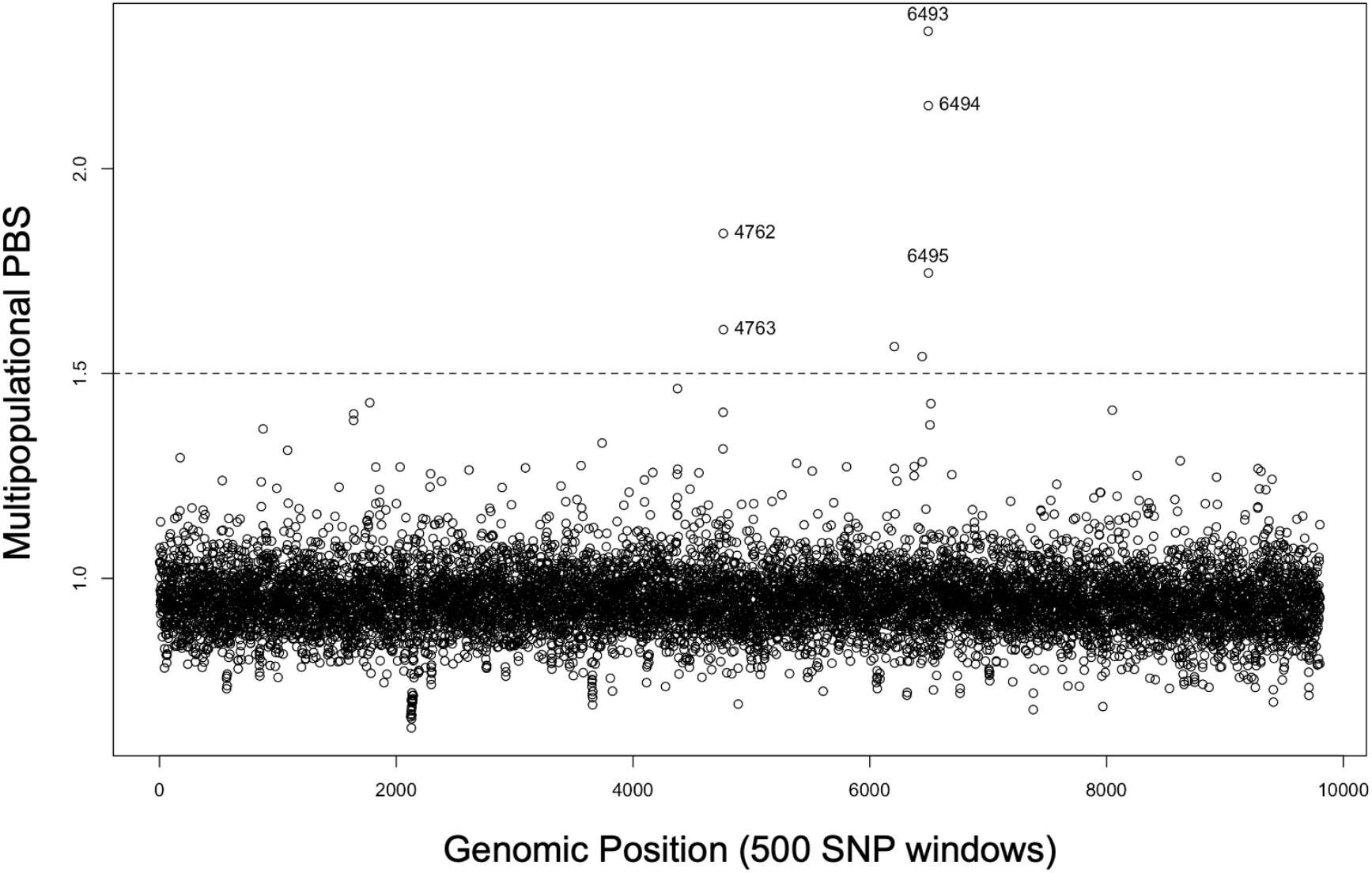
Multipopulational Population Branch Statistics (M-PBS) showing two regions in the tapeworm genome with divergent selection associated to fibrosis in sticklebacks. There were nine SNPs with relatively M-PBS (on SNP windows #4762 and #6493, see table 3) meaning their F_ST_ were much higher in HL Fibrosis Lake comparisons. This graph grouped SNPs in 500 SNP windows by 500 SNP step sizes to aid on easier visualization. The tapeworm chromosomes are not shown in this Manhattan plot due to the poor annotation of the parasite’s genome and Chromosomes positions for each scaffold are not certain. The horizontal dash line indicates the arbitrary cutoff point of 1.5 in the M-PBS; we considered the ratios above this point to warrant further analyses.

**Figure 5:**
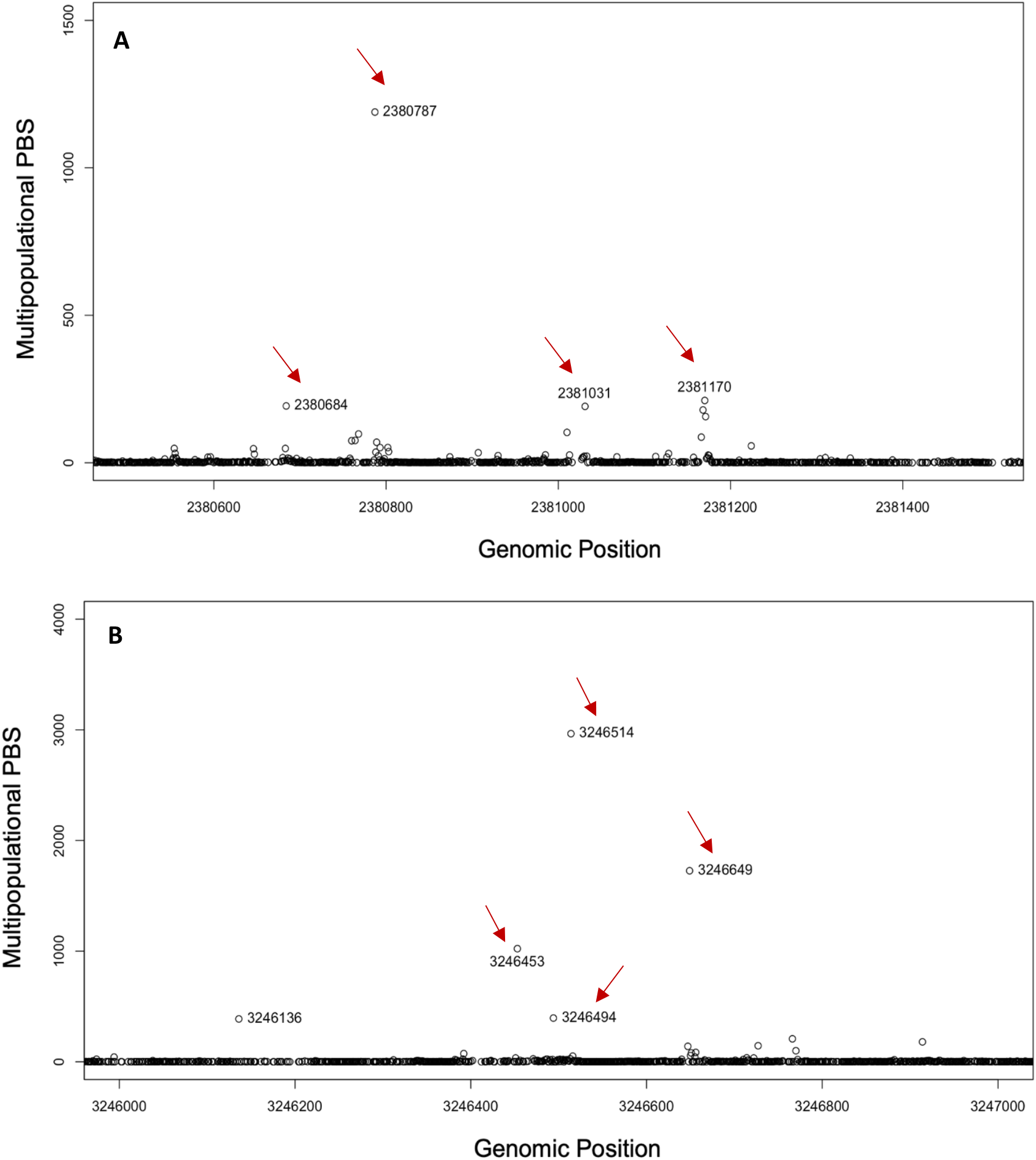
Zoomed-in graphs of M-PBS window blocks #4762 (for **A**) and #6493 (for **B**) from Figure 4 indicating SNPs with high F_ST_ in the HL Fibrosis Lake comparisons. Upon closer inspection of these SNPs (table 3.3), only those indicated by red arrows had consistently high F_ST_ across all pairwise comparisons in the HL and mostly zero F_ST_ in the HH & LL Fibrosis Lake comparisons; thus, these were kept for further analyses. For the F_ST_ values of all these SNPs, see table 3.

**Table 3:** All SNPs found in figures 4 and 5 and from M-PBS analysis. These SNPs have consistently high F_ST_ across all pairwise comparisons in the High vs Low (HL) Fibrosis Lake comparisons and mostly zero F_ST_ in all pairwise comparisons in the High vs High (HH) and Low vs Low (LL) Fibrosis Lake comparisons, except for the last SNP (#3246136) from window 6493 (see below). All SNPs except #3246136 were kept for further analyses.

| SNP window #<sup>3</sup> | SNP# | Scaffold | SNP position in the scaffold | Hight vs Low Fibrosis $F_{ST}$<sup>1</sup> | | | | | | | | | High vs High and Low vs Low Fibrosis $F_{ST}$<sup>2</sup> | | | | | | P value<sup>4</sup> |
| --- | --- | --- | --- | --- | --- | --- | --- | --- | --- | --- | --- | --- | --- | --- | --- | --- | --- | --- | --- |
|  |  |  |  | Boot vs Echo | Boot vs Gosling | Boot vs Pachena | Echo vs Frederick | Echo vs Lawson | Frederick vs Gosling | Frederick vs Pachena | Gosling vs Lawson | Lawson vs Pachena | Boot vs Frederick | Boot vs Lawson | Echo vs Gosling | Echo vs Pachena | Frederick vs Lawson | Gosling vs Pachena |  |
| 4762 | 2381170 | ScsolUL_v1_4721 | 73279 | 0.239 | 0.275 | 0.101 | 0.014 | 0.070 | 0.054 | 0.000 | 0.109 | 0.000 | 0.003 | 0.000 | 0.000 | 0.000 | 0.000 | 0.000 | **0.021** |
|  | 2381031 | ScsolUL_v1_4721 | 69642 | 0.086 | 0.100 | 0.183 | 0.110 | 0.000 | 0.123 | 0.206 | 0.000 | 0.051 | 0.000 | 0.000 | 0.000 | 0.000 | 0.003 | 0.000 | **0.004** |
|  | 2380787 | ScsolUL_v1_4721 | 66424 | 0.105 | 0.001 | 0.000 | 0.248 | 0.098 | 0.085 | 0.062 | 0.000 | 0.000 | 0.000 | 0.000 | 0.000 | 0.000 | 0.000 | 0.000 | **0.041** |
|  | 2380684 | ScsolUL_v1_4721 | 65580 | 0.123 | 0.027 | 0.000 | 0.091 | 0.117 | 0.011 | 0.000 | 0.023 | 0.000 | 0.000 | 0.000 | 0.000 | 0.001 | 0.000 | 0.000 | **0.036** |
| 6493 | 3246649 | ScsolUL_v1_8059 | 107285 | 0.367 | 0.214 | 0.164 | 0.190 | 0.460 | 0.056 | 0.042 | 0.302 | 0.239 | 0.000 | 0.000 | 0.000 | 0.000 | 0.001 | 0.000 | **0.001** |
|  | 3246514 | ScsolUL_v1_8059 | 101496 | 0.475 | 0.372 | 0.240 | 0.470 | 0.666 | 0.364 | 0.230 | 0.587 | 0.427 | 0.000 | 0.000 | 0.000 | 0.001 | 0.000 | 0.000 | **0.000** |
|  | 3246494 | ScsolUL_v1_8059 | 101434 | 0.262 | 0.019 | 0.152 | 0.353 | 0.478 | 0.091 | 0.229 | 0.218 | 0.337 | 0.000 | 0.004 | 0.000 | 0.000 | 0.000 | 0.000 | **0.001** |
|  | 3246453 | ScsolUL_v1_8059 | 100949 | 0.398 | 0.112 | 0.236 | 0.227 | 0.352 | 0.008 | 0.116 | 0.081 | 0.201 | 0.000 | 0.000 | 0.001 | 0.000 | 0.000 | 0.000 | **0.002** |
|  | 3246136 | ScsolUL_v1_8059 | 75740 | 0.000 | 0.000 | 0.000 | 0.000 | 0.000 | 0.027 | 0.000 | 0.008 | 0.000 | 0.000 | 0.000 | 0.000 | 0.000 | 0.000 | 0.000 | 0.230 |

<sup>1</sup> $F_{ST}$ from High vs Low (HL) Fibrosis Lake comparisons
<sup>2</sup> $F_{ST}$ from High vs High (HH) & Low vs Low (LL) Fibrosis Lake comparisons
<sup>3</sup> SNP window block that had high M-PBS (see figure 4).
<sup>4</sup> P values are from t-tests between $F_{ST}$ of HL vs HH & LL Fibrosis lake comparisons. Those in bold are significant based on alpha value of 0.05.

The observed peak M-PBS values are higher than null expectations, which we obtained by permuting lakes’ fibrosis status. There were only five possible permutations that retained the geographic structure and three populations of each fibrosis level (see supplementary table 3). We used three different test statistics to see if the ratios from the original observed scenario was different from the other random scenarios (i.e. Max M-PBS ratio, number of points exceeding M-PBS ratio 1.5, and mean of the five top M-PBS ratio points). The original observed scenario consistently had higher test statistic scores (see figure 6, supplementary table 3, and supplementary figure 1). For instance, we found that the observed scenario has higher average ratios when averaging the top five ratio points of the M-PBS (figure 6 and supplementary table 3). The observed scenario also has higher maximum M-PBS ratio and higher number of ratios exceeding M-PBS 1.5 (supplementary table 3 and supplementary figure 1).

**Figure 6:**
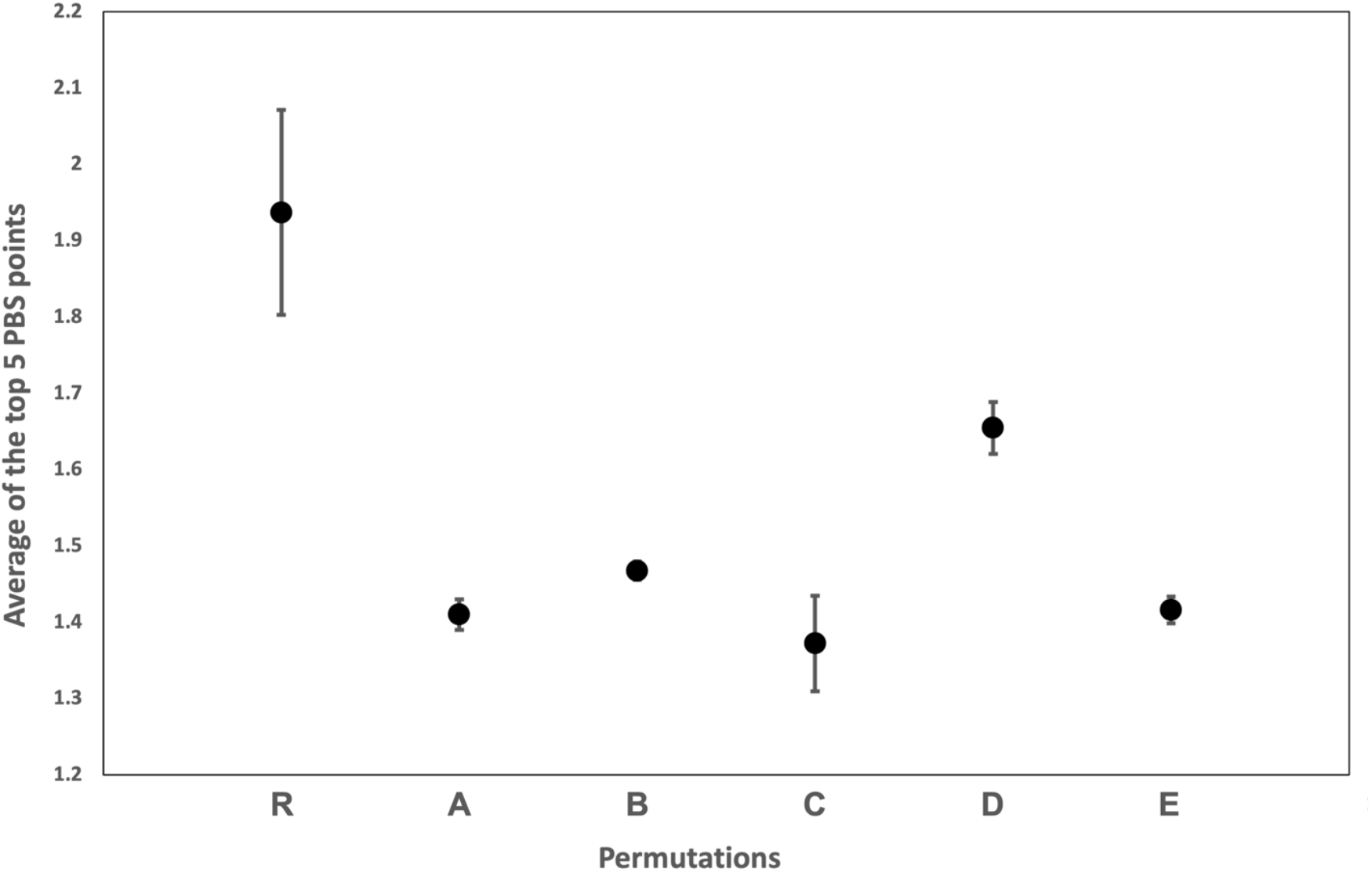
Average of the top five points of the M-PBS for all possible permutations of high and low fibrosis lakes indicates that the real observed scenario (i.e. R in the x-axis) has the highest average (see supplementary table 3 for all permutations in fibrosis scenarios).

In case the F_ST_ values were due to lake-specific duplicated SNPs (i.e. duplicated in some lake populations but not duplicated in the reference genome), we also run correlation tests between SNP read count and F_ST_ and M-PBS values. If the F_ST_ peaks seen in the M-PBS were due to duplicated SNPs in some lakes, then there would be a significant positive correlation between read count and F_ST_ and M-PBS across the entire genome. Results of correlation tests indicate that there was a significant but negative correlation between read count and F_ST_ (t = -177.07, df = 3677268, p-value < 2.2e-16, correlation: -0.092) and M-PBS (t = -170.05, df = 3677268, p-value < 2.2e-16, correlation: -0.088). This indicates that the F_ST_ and M-PBS values seen was not due to lake-specific duplications in the tapeworm genome.

We extracted 1.2 to 1.4 kb sequences enclosing the SNPs in table 3 (except for SNP #3246136 from window # 6493) in FASTA format using IGV version 2.11.1 (Robinson et al., 2011) and performed a BLAST of these sequences in WormBase Parasite (a repository for parasitic worm genomes; Howe et al. 2017) and in the National Center for Biotechnology Information (U.S. National Library of Medicine, 2022) genome repository websites but found no known genes (or genes to known functions) related to these sequences from the sequenced *S. solidus* genome nor from any other organism’s genome. See supplementary notes for more details on these sequences and the BLAST results.

## Discussion

The tapeworm populations in this work are not very divergent, even for those that are ∼130km apart. This can be noted in the genome-wide F_ST_ in all the population comparisons, which is generally low on average (table 2). However, there are a few targets of divergent selection among population pairs (figures 2 and 3 and supplementary table 2). The SNPs from these targets of selection seem to be related to local adaptation as they are found mostly in comparisons between lakes that are geographically distant from each other (supplementary table 2).

Furthermore, through the M-PBS analyses we found at least eight SNPs from two genomic regions in the tapeworm that were associated to a phenotype involved in parasite resistance (i.e. fibrosis) in sticklebacks (table 3). Fibrosis is one of the main stickleback defense mechanisms against tapeworm infections (Weber et al. 2021), and as such it should act as a strong selecting agent in the parasite. Weber et al. (2021) discovered several candidate genes involved in fibrosis expression in the fish; these candidate genes were determined through laboratory infections with the tapeworm. The SNPs found in this work might be associated to -or be part of- genes in the tapeworm evolved to adapt and overcome fibrosis in fish, and as such these SNPs (and their associated genes) might be coevolving with the stickleback fibrosis genes from Weber et al. (2021); however, association studies with more fish and tapeworm populations will be needed to validate this premise. Note also, that besides these tapeworm SNPs could be an adaptation to the expression of the fish fibrosis genotype, the fish fibrosis might also be an indirect genetic effect of the parasite genotype (e.g., different alleles in the tapeworm might be the cause of some fibrosis differences among lakes; for instance, Hund et al. [2020] has shown that local tapeworms caused more fibrosis in local lake fish, compared to tapeworms from elsewhere).

Nuismer et al. (2017) highlighted the importance of host-parasite coevolution in driving important evolutionary transitions such as asexual to sexual reproduction and haploidy to diploidy, and how genetic studies of coevolution can offer rich avenues of research in evolution. They also highlighted the current difficulties in studying the genetic basis of host-parasite coevolution due to the of lack of known interacting genes in the hosts and their parasites. Although the SNPs we identified here has not been proven to interact with genes responsible for upregulating and downregulating fibrosis in sticklebacks from Weber et al. (2021), there is the potential that these genes are associated and can offer future researchers important tools to study the long term dynamics of coevolution at the genetic level.

The SNPs (and their prospective genes) found in this work are also potential tools to study how tapeworms escape hosts defenses at the genetic level, which will have important implications in immunology and animal husbandry. Helminths, including tapeworms, have a detrimental effect in human health and agricultural output, especially in developing nations (Weber et al. 2017), thus, understanding the molecular mechanisms of how these parasites elude hosts defenses will be indispensable in the developing of drugs and strategies to combat infections, and having SNPs of possible genes involved in parasite escape of host defenses is the first step in this process.

However, after performing BLAST searches in two genomic databases (WormBase and NCIB), we did not find any known genes (either from other helminths or from well known model systems) matching the DNA sequences containing the SNPs found in this work. This is hardly a surprise given the poor annotation of the *S. solidus* and other helminth genomes. Thus, better annotation of the tapeworm genome followed by further experimental studies (e.g. gene knock out experiments) will be needed to elucidate the function of these genes and how they are involved in adaptation to fibrosis in sticklebacks.

As mentioned before, we also found SNPs from genes that might be related to tapeworm local adaptation. Pachena and Frederick lakes have several SNPs with high Fst (0.72 to 1) when compared to lakes from the Campbell River watershed (see figures 2 and 3 and supplementary table 2). These two lakes are separated to the Campbell River area by ∼130km (see figure 1A), so it is very likely that these SNPs are associated to local adaptation. Besides Pachena and Frederick lakes, Echo Lake also has five SNPs with high Fst (0.74 to 0.79; see figures 2 and 3 and supplementary table 2) to the other lakes of the same Campbell River watershed. Again, this might be due to local adaptation since Echo Lake is further apart (>10km, figure 1A) from the other Campbell River Watershed lakes used for this work (i.e. Boot, Gosling, and Lawson). Local watershed temperature and water chemistry might cause these local adaptations since the complex life cycle of the tapeworm involves deposition of its eggs in lakes for an unknown period of time and a brief free living larval stage (Dubinia 1980). A more thrilling possibility is that these local adaptations are due to differences in host populations (i.e. either copepods, sticklebacks, and/or piscivorous bird populations, all of which are hosts to the tapeworm [Dubinina 1980]). In the case of sticklebacks, phenotype(s) other than fibrosis might be acting to produce this tapeworm local adaptation. Future reciprocal infections are necessary to confirm tapeworm local adaptation due to hosts.

Nuismer et al. (2017) also provided a statistical framework to identify genes involved in host-parasite coevolution through covariation of genes across space (i.e. local adaptation) in the species involved. This method works best for host-parasite systems where local adaptation has been shown to be strong. The extremely high F_ST_ (0.7 to 1, supplementary table 2) in the SNPs related to local adaption found in this work is a robust indication of strong locally adapted tapeworms. Thus, it might be possible to use the stickleback-*S. solidus* model to test the Nuismer et al. (2017) methodology to find more genes involved in coevolution; this is of course, again after further reciprocal infection studies between tapeworms and fish populations from the two (and probably more) regions.

After over half a century of intense use (Dubinina,1980), the stickleback-*S. solidus* tapeworm model is still proving to be a rich system for host-parasite research, and with the advent of modern high throughput in genomic sequencing and immunological assays, coupled with big data analysis methodologies, this model system should provide even more fruitful research in the decades to come.

## Supplementary material

**Supplementary table 1.** DNA quantity per tapeworm from each pooled lake:

| lake | n individuals<sup>1</sup> | ng per ind added<sup>2</sup> | qubit ng per ul<sup>3</sup> |
| --- | --- | --- | --- |
| Lawson | 21 | 27 | 6.58 |
| Frederick | 24 | 30 | 2.72 |
| Boot | 23 | 30 | 3.35 |
| Pachena | 23 | 48 | 10.85 |
| Echo | 25 | 50 | 16.10 |
| Gosling | 24 | 30 | 7.13 |

<sup>1</sup> The number of individuals present in each pool.
<sup>2</sup> The average amount of DNA (in ng) of each individual that was added to a pool. This varies, as the amount of DNA extracted varied between individuals in each population.
<sup>3</sup> The DNA concentration of each pool in ng/uL.

**Supplementary table 2.**
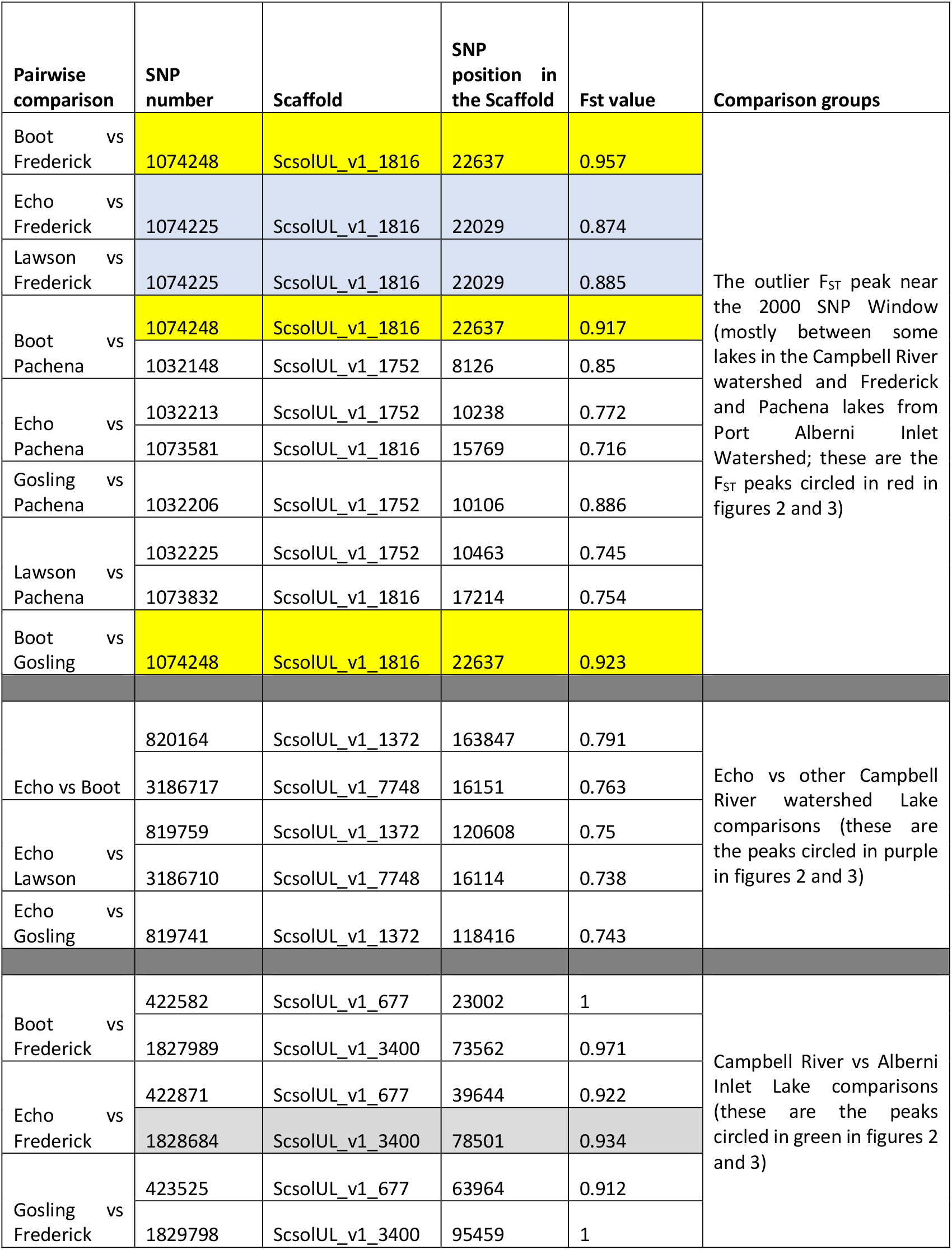
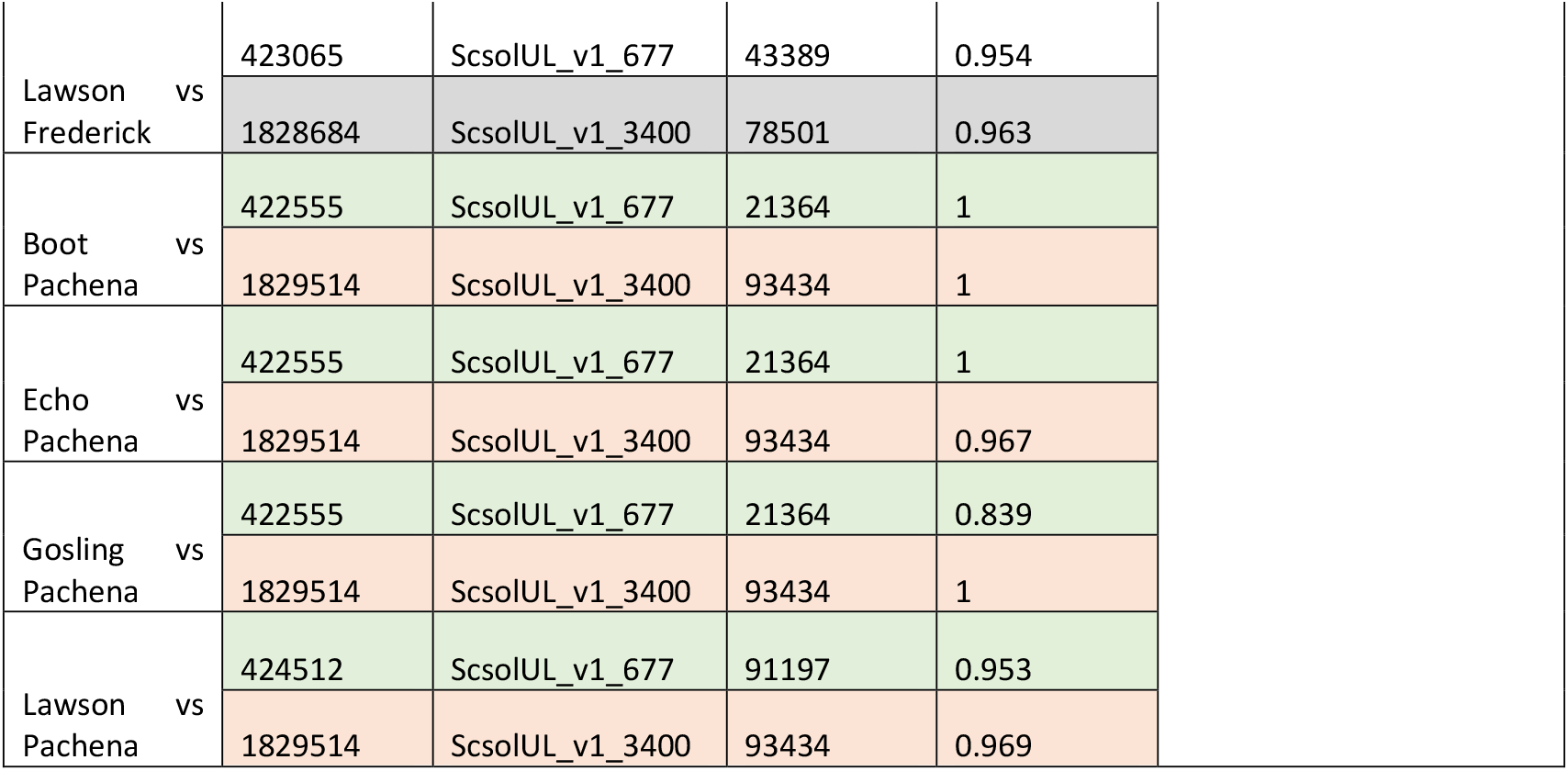
Local adaptation SNPs found in the Manhattan plots of F_ST_ pairwise comparisons (from graphs in figures 2 and 3). Rows highlighted by the same color indicate same SNPs.

**Supplementary table 3:**
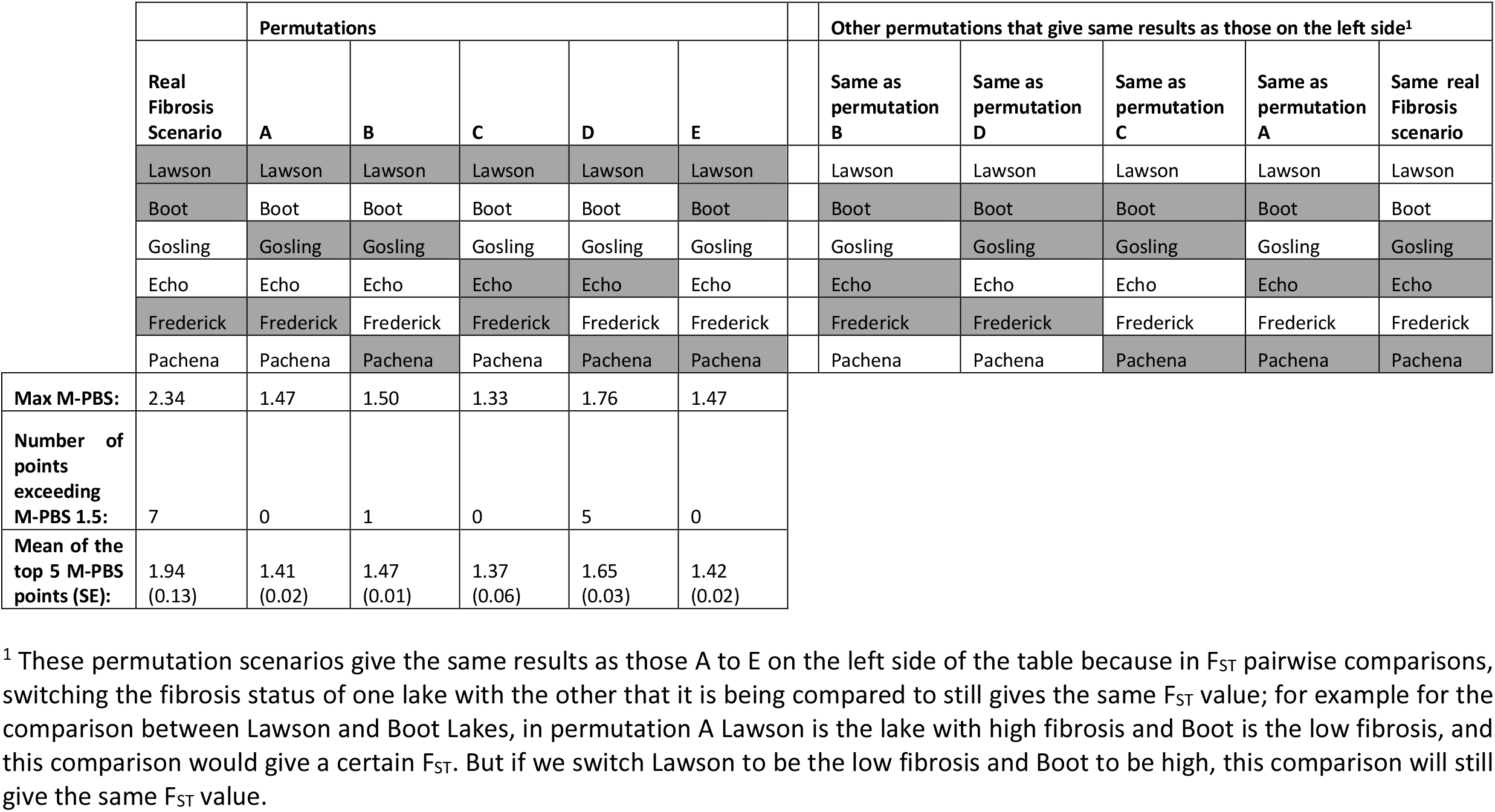
Testing different fibrosis permutations by randomly assigning high and low fibrosis status to the lakes. The real fibrosis scenario still had higher maximum M-PBS ratio, higher number of points exceeding M-PBS 1.5, and higher mean of the top five M-PBS points than the other permutations (see figure 6 and supplementary figure 1 for graphical representations). Shaded cells in grey indicate high fibrosis lakes; clear cells indicate low fibrosis lakes.

**Supplementary table 4:** average number of reads per site per population pool.

| Lake (population pool) | Average number of reads |
| --- | --- |
| Boot Lake | 36.08 |
| Echo Lake | 36.76 |
| Frederick Lake | 29.61 |
| Gosling Lake | 29.84 |
| Lawson Lake | 38.89 |
| Pachena Lake | 46.39 |

**Supplementary figure 1:**
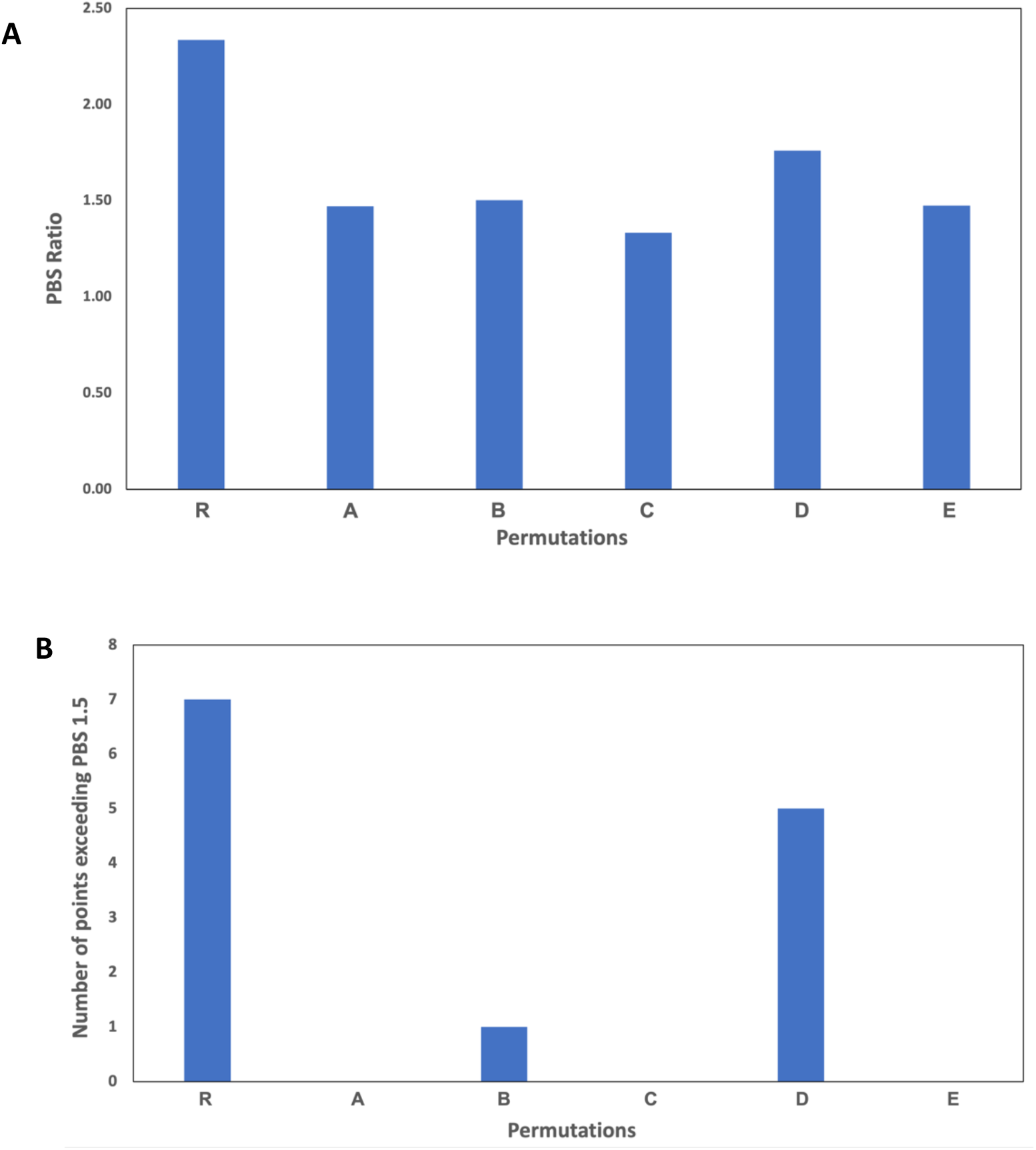
the real case scenario (i.e. R) still provides the highest maximum M-PBS value (panel **A**) and the most number of points exceeding M-PBS 1.5 (panel **B**) from all possible permutations on the high and low fibrosis lakes. See Supplementary table 3 for all the permutation scenarios.

### FASTA sequences adjacent to the SNPs from table 3 and their BLAST results

ScsolUL_v1_4721:72679-73879 (1200bp, SNP of interest #2381170 from SNP window #4762, which is in the middle of this consensus sequence from IGV_2.11.1) GTCTGATAGTAAGGATAGAGGATAAGACCGCAACGGACGCTGAGAACGTACCTCGCCTGCGCAGCTCAGTGACCGTAGGGGAGTTGAGGGAATGCGGGACTTTAGTTGGCGGCTGCATCATCTCTTCCATAC

GATCCAGGGAAATGGAGGAGAGTGAACGCGTGACAGATTTGGGTCTTCCAGRCGCAGAGGKGCAGCC GGAAGGTGGAGGAAGAAGAGGCAACGCCGAAATTCGCAGTCGTTGTGAGGATTGTGCTGGTTGAGTCACGTTGGCTAGCTGCAGATAAAAAAATTAAATGGCGGAGTTGCGATAACAGCATTGAGGTCCAAAACAGGAAGCAAAACCTGAGTAAACTTAATGAGGACCCTTGTTGGATGACAGTCTCGTCCAGAGGAGAGGTG GAYAGTGCAAGAGGCAAGAGGTCATCGTTTAAACGGAGGGGAGCTGCAAATCGGGGTTTATCGCTTGCAGTTTCCACCAYACACGATGTCCAAGCAGGCTAAGAAGGGTATAAACGGGGAGATGAGGGGCAAAGACAGAAAGAGAGCAGACTGGTGGAATCAAGGGGRCCCCRSCAACGCCCAAKAAAWTTCYRAGWTWKYTTACATTGGCTCRTCMGACTGAAGTGMCATTGCAGTGCCTCMCTTGTCCGCRGCATCAAGCAGTCCCGAT TCAGCTCACTAGTCAACTAATTGCCCGTTCCAAAACGCAGTCAGAGACAAACGCACCGACACGGGTGCAACTCACGCAACGTGCGCCTTTTCGTTGTGACCCATAAGAAAATTGTCAACTCACTGCAATTTCTCATTGTTCTTTGACTTAGGTTACTGTGTGCGGTCATAGCTACACAGAAATTATTCACTGTCACGGCCCCACAGGCAGAGNNNNNNNNNNNNNNNNNNNNNNNNNNNNNNNNNNNNNNNNNNNNNNNNNNNNNNNNNN NNNNNNNNNNNNNNNNNNNNNNNNNNNNNNNNTTGGTCCATGAACTCGCCCCCTCKTCTTCTTCGT CTTCTTCAGGAGATGATGAACGCGATATATATATGAGAAGAAAAGGCGATCTCAGGGACTATAGGTGTATTGTCCTGACGTTTCAAAAGCATACAAGCTTCCATCGTCGGAGGTGAGTATCATACCGCGTCTCACAGTATTTATAGGGGGCGATAGCCATTTTTGTAACACAGCGTTGATTGGACGGGTTTA

Note: This sequence BLAST-ed to a novel unknown protein coding sequence named SSLN_0001547401-mRNA-1 in the WormBase Parasite website.

When BLAST-ing it in the NCBI website, it primarily BLAST-ed to the parasitic tapeworm *Spirometra erinaceieuropaei* scaffold SPER_scaffold0052000, using both the megablast and BLASTn search engines, with E value of 4e-40, 83.78% identity, and 15% query cover (for megablast).

When BLAST-ed to NCBI’s Transcriptome Shotgun Assembly (TSA) database using the BLASTn search engine, it matched primarily to *Schistocephalus solidus’* ssol_TR151174_c0_g1_i2 transcribed RNA sequence (sequence id: GEEE01020411.1) and ssol_TR151174_c0_g1_i1 transcribed RNA sequence (Sequence ID: GEEE01019478.1), both with E value of 8e-103, 97.35% identity, and 30% query cover. Both codes for an unknow gene.

ScsolUL_v1_4721:69042-70242 (1200bp, SNP of interest #2381031 from SNP window #4762, which is in the middle of this consensus sequence from IGV_2.11.1)

TTTGTTKGAGAGTTCGAGCACAGGCGSGGTCCCAGTTARGATTGGTCGTTCGAGGTTCTCTTCG CTACCACGTTGGATCAAGGCCTAACCTTATGAGCTCAGTGTTGAAGCCCTCGAGCCGGCGCCACTTAAA AGGAAGTAAACTCAAACCCTGGCTTGACACAGWTCGTCATGRYATGGAAATGGTTCTYGGAATKTGGGTCGAGATCGCCMAAWCCRTTCTAATAATTKTTGRTCTGGCATAATTCCRGTCAATAGCTTGTTYATTTGAACTTCTAAAGATCTCTGAAAGATTTGGCCTCGAWAATATCCCCCCAATCTTGGCTACTTTCTCGGACCCCTC TAATTTGCATAGAAAGCCAGAAACAGTACCTGAAACACTGTTGCCCGTGTGACCATCTCCACTCTTACAAACCTAGACAAAGGTTATGGTGCATGCAAATAGCCACGTATTCGTCCGTTTATTTACTATGTTACTCGTGAACTATCGAACATAGTGCGCATTCTATATCAGATCATCAAAAACACAGAAAAATGTATGATTATCCTTTCATGGAAATTATACATGTGTAAATTTCATCTGTCTGCAATTGTTGGATTCTAAACTTAGAACGCATATGCAACCACAGAAGTAGTCTCAATATTATTTAGGAATTGCAAAAGTTTCTAAAATTCTAAAGCTACCCTTTCTTCGGGTGTTAAAATAAGACGTATTGCTACCAAATAATTACAATAAAGGATATGTGAGCACCCCGCTCGAACATATTCCAGTTTATCAAGGCATGGAAAATCAGCGAAAAAACTCATAATAAAAAGAAGACATTGTTTTATCCAGTAGTTTCTTCAGATGTCTGAAAAGAACTTCATTTAAAAGTCAGCATTCTGGCGAGCATGAAACATCTCAGG ATCAYTMTGCATGGTTATCAATATTAAAAATCCACCTAAATAATCTTTCCTAATCGAAAACGGACWTTAR GGAAATTCGGTTAGCATTTCATAAGATTGGTCATTGCCTGTACTAACGATACTGCTGATTCGGACGTTCT TCAAMCCGTTAAAAAATAGGCTGATGYCCAACTTCTTAAGGAAAGTTTCGATCAATCCTCAGCGTGACTTGRCAGTCATATATTTAGCAGTCCRATTAAGTCGTTCATACGGACGKGGAATGCRAACGTACAARCTAACCTTYTTTGAGWAWRGG

Note: This sequence BLAST-ed to Scaffold SSLN_scaffold0007263 in the WormBase Parasite website. We should note that an unknow gene from transcript SSLN_0001547401-mRNA-1 was located just before the sequence above. This gene is the same one from BLAST result from SNP #2381170 earlier.

In NCBI, this sequence BLAST-ed only to the parasitic tapeworm *Spirometra erinaceieuropaei* scaffold SPER_scaffold0017214 when using the megablast option (with only 6% Query cover, E value of 4 e-05 and 89.49% identity). With the BLASTn search engine, it BLAST-ed primarily to *S. erinaceieuropaei* genome scaffold SPER_scaffold0063623 with E value of 4e-28, 73.86% identity and 21% query cover. Both do not contain any known genes.

When BLAST-ed to NCBI’s Transcriptome Shotgun Assembly (TSA) database using BLASTn, it matched primarily to *Schistocephalus solidus* ssol_TR155804_c10_g2_i3 transcribed RNA sequence (sequence ID: GEEE01017489.1) which codes for an unknow gene. But this match only had 7% query cover, with an E value of 5e-10 and 78.26% identity.

ScsolUL_v1_4721:65280-66724 (1444bp, SNPs of interest #2380787 and #2380684 from SNP window #4762 are enclosed in this consensus sequence from IGV_2.11.1)

GCGGTACACGGGTTTTTATGTGAAATTAATGCCCTTTTTTCTGTTAAGCCATGCTCAACAGTCACCATACACGCAATTCTAATAACATCCAAATAACGTACTCATTTGAAAAACGAAGCATTGTTTTACCAAATAAATACCTGCGGCGGTGAAGAGATGCTTGCCCTCGCTGGCAGCCTCAAAAGCAGAAAGAGCAATGCCAACGTTATGAATCTTCTGCAGACGGCTTATAGGTGGAAACCGGACAAGACTCATCAAATTTCCAACATGTGAT AAAATGGAGCCAGAATCGTTTAGCYCTGAATGATCCGTTGACACTTTTCCTGCGAGAAGCTGGTCAGCCAATTTTCTACAAAGAGAAACCCGACAAGATTACTGTGCGGTTTGGGKTGCAATGAGAAAAGCAGCGCGGAACGGAATGCATAGCTCACACGAGACGTAGACCATCGCGCAGATCTACGGCAAGGTTCTTCASAGTGA AATCAAATTCGTCAAGGGCATTTTGCGAGACGCAGACGGTGGCTCCGATGGCTGATAGAGCTCGCACTAGGTTAACTTCACCCTGGAGACAGAGGTTGGCGAACTTCAACAAGACCTCGGCAGAGCTCTGGAGGCGA AAGGGTAGCAGTTAGGACTTGCAAACAATGCAAATTCGATAGTTTTCACAGCGATAGTATGCACGCACCTTTATAATTGACTTTGATTTGAAAAGACAAGGGTTGTAACGAACAACCTGCTTCGCCCTCGACTGATCCAGGAACCAAATTAGGACAAGACAACGCTTTACGATCTGACAGTTGTAGTGATCGTCGAAAGTGACTTTAA AAATTWAAAATAAAYAAWTTAAATGGAGACACTAACYARTGAGAAAGAGGGCAACATYTYGGTTGAGGTGCATAAATAAAAAAAATACCTGGGACATGTTGATTTCGTCCCCAACYGACCGACTTTCGAGCGTTCGGA TGTTTTCCCTTAATGCTAGGTTCTTCTTCTTTCCACTTGGCTTTTGGCATTGGAGGTACGGCACCGAAGATGTAAGACTGAACACGACCGGAAGTCGTCAGCGATTTCTCGCAGGCGTTTACCGTGGTGGGAGTGGGAGCTGTGAAGGCAGACAGAACGTCCACACAAAGCTGTAGCCATACGGGCGAGTAATGTGCCAGAAAGATTTCACAAATTCGATTTTGAAGACCTGCAGAAAGAAAGGAATGCTCAGGGTTAGCTCTATTTGAAACTTGCGAATTGACTTACATGACGAAGGTAGGAAAACGTTTAGGTATTTATGATCCCCAAGGGTTCCCCTCCTCCC AAACGCAGTCCARAATAAASAGTAMAAWAAAKCCAATWAAGGCCTACACGTAAAACAGTTAGCATACGAGAGTACAGGATGRCCTGTTCCAAGTGAAGAAAACCAGTACGCAGATGATGAACTGACGATCTCGGG G

Note: This sequence BLAST-ed primarily to a novel unknown protein coding sequence named SSLN_0001547401-mRNA-1 in WormBase Parasite website with E value of 7.9e-121, 81.2% identity, and only 9% query cover. The same match result was found for SNP #2381170 earlier. No significant match similarities found when using megablast search engine for NCBI. With the BLASTn search engine, it BLAST-ed primarily to *S. erinaceieuropaei* genome scaffold SPER_scaffold0024198 with E value of 2e-116, 77.62% identity and 56% query cover. This scaffold does not contain a known gene.

When BLAST-ed to NCBI’s Transcriptome Shotgun Assembly (TSA) database using BLASTn, it matched primarily to *Schistocephalus solidus* ssol_TR151174_c0_g1_i1 transcribed RNA sequence (sequence id: GEEE01019478.1) with E value of 0.0, 94.49% identity, and 61% query cover. Its second highest match was also to *Schistocephalus solidus* ssol_TR151174_c0_g1_i2 transcribed RNA sequence (sequence id: GEEE01020411.1) with E value of 3e-103, 90.39% identity, and 55% query cover. Again, these results are similar to SNP #2381170 above. Both sequences codes for an unknow gene.

ScsolUL_v1_8059:106586-107981 (1395bp, SNP of interest #3246649 from SNP window #6493, which is in the middle of this consensus sequence from IGV_2.11.1) TCSTGCAGTAGTTTACTTTTGTTGCTTCCTGTTTCGTTATATTTGTGGACCCTTTATTTCTTATGTTTTTTTGATAGTTTTTTTTTATTTGTGCACCTTTTTGTATNAAATMTTTTACATTCCAACCCCCTCGGACTTCG AACCCTGCCCCACGGGTCACTATCGTTAAGCAGAAACTTTAATAATGTATGCACGAAGGACTACCGATAAAGACAAAGGGGCTGAACTGGCCGTTTCTGACTTATTTGTCGTCGTCAGGCAGCCCTACCCAACCCCAGGTTTGGATATCTTGTGATAACGTTTATTGGTACTCCTTCCTATGTATATTACGGCTCACTAAGCGATTGGCTGCAATAGATCGGGCTTGAAACCCAAGACCCAATACTATGAGCATGTGCACGTAACCCGGTCAATGAACGCTTTTTCTACGGTAGAACATTTAGTGGCTTGATGAAGTAGCGCTCACCGATGAAGCTACGGAACATGCTTGTCCTTTTGTGCCTTGGGGCTGAATCTAGAGCTATCACATGCAGTTGCATAGCGACTAGTAACATATACG TATGATGGAGTACCGACAAGCACAAGGGACRCCARAATGGCCATTTCTGSCTTWTCAGCMGCCATCAGCCAGTCATCCCCAATCCAAAGTGTAGCCCACCTGGGATTCGAACCGTCGACCTTTAGTCATTGAGTGRAA CGTCCAACCAATGCGCCACGGGCTAGTTGTCTTGAGCCCCAAAGCACAACCTGACAAGCATGTTCCTTAACTTGATCGGTGAGCGCCACTTCGTCATTGCTATTCCCGATCACAACCGACAGGACAACGAGCCCGTGGC GCATTGGTAGAGCGTTCGACTCAATTACCACTTGRGATTGTGTATGACTGGCGAATGGCGGCTAATAAACCAGAAATGATCACTCCGGTCTCCCCTTTCTTTGTCGGGTACTGATACTCCATCATTTAKTGGTTTCCYAGCTGTCCAGTAGAGCCAGTTCGCSTTGTATATSTCATTCACTYGCACCTTTTTCTCGTTTTAGCGATCACGACG ATCCGAAGCGCCCGTTCAATCGACCGCCTCCGTAGCGAACACTCCCCTTACCCATAGGGGCGTACGTGT GCGTGCCTCYATGACGGTGAGTCATGTGGGGCTTGAGTGCGTTTTTTTGTATTTGCCTCACTTCACTGCCC GCTGTTTTAACCTACATATCTTCCTGTGCTTTAGACTTAATGACAGGACCAACAAGGAAGATAACTGAATAAAGCTGTTTAACCTACTTAACTGGAACTGGTGTACGAAGATTTTAGTTAGTGTCTGACGGGTGACAGA GGGTGGGGTACTCAGTGGGGACTYGTACATGGTWGGAATTTTARCCAGGGACACSGTCGACCCAGAAT

Note: The above sequence BLAST-ed to an unknown protein (probably obtained from mRNA sequencing) at WormBase Parasite website. The id of this transcript is SSLN_0001651701-mRNA-1. Searching this transcript in the NCBI website provided no description of what this gene might be or do. After BLAST-ing the above sequence in NCBI’s website, we got matches to the tapeworm *Spirometra erinaceieuropaei* scaffold SPER_contig0050188 but not to known genes.

ScsolUL_v1_8059:100436-101833 (1397bp, SNPs of interest #3246453, #3246494, and #3246514 from SNP window #6493 are enclosed in this consensus sequence from IGV_2.11.1) ACGTCSTTTGTGCTTTAAATTTCCACGCTATTTATTTGCCATGGTCTTACGTTAAGCTCTACCTTTATGCATATTGTGCCCCCTCCCACCTCGTACGTCTTATTTCATGMCTTTTGAYWAAGAATTCAGCTCACGAGCTAAGTGTACKGTTTYKTCAMCTTATATTGTATTTGRGCATTCGTTASTTACRCYGCCAACTTCAACTCAATTACCTACCTATTCATAGATAATCACAACACCCACATGCATGTTATMCAACAAATGTCCTAATTTTATAGGA AATCAAGATATCCGAAAGAAAGATGAGCCTTTTATGACGTAACGTCACCAAATTCTGTTTTCCTCCTCTTCTTGATGTCTCCGCAAGTTTGTCCATTTTGCCATATTATTTATCTAAGTATGCTTTAAAAATCATCACGCCTA AAGAATATAATATATTTTATTGTKTTTTGMCACTGYCAAGGTTCKTSAYTCCTCTKAAGCYTCAMCGCAGTCTTCCGYCGTCTCCATTCGRCTGCAGGCAGGAGTGCCTTTTCAGGGCTCAAGAACTSTTAGTKTTACTGA GTATTCTTGTCCGCCCTCCTGTTAGTGTAAATTTCATTTCCCGTCTGTTTTCGTTGCTGCGCGATGTGGCTCATTTTTTACTCAAACCTAAAGCGTTTGCGCGAGCCGATAGAGAATGTCGAATGGGTAACCTGTGGCATG GATAGGGTCTGTGAACATGCTGTTCTGAGCAGACTTGAGCGARAGCGTCMTCGTTGGAAGTAGGCTATGAAATCTGGCGCATYAACTGATTGYTTCGTCCAGCRGGGTGTYCAGTGATGTYKTTTTWATCCGTGWWCTCTTYAACTATCCCTCTCTARCAGTRGGGRTGCTCGATMACTTCTGAYTGAGGACCATCCTTTGGGTGAAGCATACCGRCTTAGTTCCGAGTGAGTCGGYCTGTACKYGCTWCCATAACATTATGAAMATATWTAATGTCAYCCAAGAAAGACGACRTAGCTAGTTTGGCTGCGTTCTTCGTAGTGCTCATCATGGAATCCGTTTGGCT GCCCTCGATTTGAAGTCCATGCCCATCTAGGAGTGTCGGTGAAGCCTTCTCAAAATCTGTCTTGACACAGGTCGTCAGAACATTGAGGTAATTCTTGGGCCTTCGGTACTTGGTATCGGTCGCCAAAGCGGGGAATGCATCTGGCTGTCCAGATCTGCTGCTCCGGATCGTACGACATCATTGAGGCTGCCTAGATCAAACCTAATCAC TGACGTACCAAGTACAAGTATTTATGACTKACGCGTGCCCTAATCGAATGCTGGTCCATACCACTTACAC TGCTGGTCTTAGCTGCTGAATGTGACACACAAACACCAAATGTCCCTGTCCCCCTGCACCTGAAATGT

Note: The above sequence BLAST-ed to a *S. solidus* genome region with not known genes (not even known RNA transcripts) at WormBase Parasite website. After BLAST-ing the above sequence in NCBI’s website, we got matches to the tapeworm *Spirometra erinaceieuropaei* scaffold SPER_contig0140472 (but barely with only 28% Query cover, 65.80% identity, and an E value of 4e-23) and not to known genes. When BLAST-ed to RNA transcriptome Shotgun Assembly (TSA) in NCBI, we found that it matched 2 reported *S. solidus* RNA transcripts: ssol_TR126982_c0_g2_i2 transcribed RNA sequence (Gene Bank accession number: GEEE01009784.1) and ssol_TR126982_c0_g2_i3 transcribed RNA sequence (Gene Bank accession number: GEEE01009770.1), but both only had a Query Cover of 6% and E value of 2e-28. Both transcriptomes are from genes of unknown function.

## Notes

### Competing Interest Statement

The authors have declared no competing interest.

